# wrmXpress: A modular package for high-throughput image analysis of parasitic and free-living worms

**DOI:** 10.1101/2022.05.18.492482

**Authors:** Nicolas J. Wheeler, Kendra J. Gallo, Elena J. Garncarz, Kaetlyn T. Ryan, John D. Chan, Mostafa Zamanian

## Abstract

Advances in high-throughput and high-content imaging technologies require concomitant development of analytical software capable of handling large datasets and generating relevant phenotypic measurements. Several tools have been developed to analyze drug response phenotypes in parasitic and free-living worms, but these are siloed and often limited to specific instrumentation, worm species, and single phenotypes. No effort has been made to unify tools for analyzing high-content phenotypic imaging data of worms and provide a platform for future extensibility. We have developed wrmXpress, a unified framework for analyzing a variety of phenotypes matched to high-content experimental assays of free-living and parasitic nematodes and flatworms. We demonstrate its utility for analyzing a suite of phenotypes, including motility, development/size, and feeding, and establish the package as a platform upon which to build future custom phenotypic modules, including those that incorporate deep learning techniques. We show that wrmXpress can serve as an analytical workhorse for anthelmintic screening efforts across schistosomes, filarial nematodes, and free-living model nematodes, and holds promise for enabling collaboration among investigators with diverse interests.

## Introduction

The past decade has seen the development of a variety of software for the acquisition and analysis of high-throughput and high-content imaging data of roundworms and flatworms, both free-living and parasitic [1,2]. New instrumentation and analytical capabilities have laid the foundation for a new era of phenotype-driven screening for anthelmintic compounds.

Early iterations of image-based screening focused on gross worm movement, using a number of different approaches to quantify movement, including sparse measures of optical flow and frame-by-frame pixel variation [3–7]. Optical flow was found to be robust to a number of diverse nematode and flatworm parasites and has been the basis for some of the largest phenotypic screening efforts to-date [8–10]. Other developments in high-content imaging, sometimes combined with the employment of fluorescent stains to reveal fine-scale phenotypes, now allow for the quantification of detailed morphological and molecular features that can be used for image-based classification strategies [11–14]. Open-source packages have been developed to more readily handle large imaging datasets and provide quick readouts for quality control of entire experiments, plates, wells, and even individual worms [15].

Not unexpectedly, individual labs often develop their pipelines to suit their own needs. These pipelines tend to focus on specific species and stages, require specific instrumentation, and demand an advanced grasp of compiled languages, resulting in siloed development and redundant rather than collaborative engineering efforts. There have been recent developments that unify parts of these efforts for the capture of phenotypes in model nematode species [15]. No package has yet to bring multiple phenotypes (i.e., motility and morphology) into a single framework that prioritizes flexibility across free-living and parasitic worms. Here, we present wrmXpress, a modular open-source package that consolidates multiple analytical approaches. It is written entirely in popular, open-source, interpreted programming languages (Python and R) and is configured with a human-readable markup language (YAML). It is containerized for deployment across a wide variety of compute platforms (both distributed and isolated), enabling collaboration and reproducibility. Finally, while it ships with a range of phenotype pipelines, it establishes a foundation for extension to additional analyses and species, including future image-based deep learning applications.

## Methods

### Protocol and data availability

wrmXpress v1.0.0 is publicly available at https://github.com/zamanianlab/wrmXpress and includes a Conda environment file to install dependencies. A Docker image that includes all dependencies is publicly available at https://hub.docker.com/r/zamanianlab/chtc-wrmxpress. Example imaging data for each module is available as a Zenodo repository (10.5281/zenodo.6558304).

### Image acquisition and analysis

Images were acquired with an ImageXpress Nano (Molecular Devices). *Caenorhabditis elegans* was imaged at 2x with transmitted light and GFP/TxRed where applicable. For *Brugia malayi* microfilariae motility, 10 frames were acquired at 4x with transmitted light with the field of view focused on the center of the well; for *B. malayi* microfilariae viability, wells were imaged at 4x with excitation at 472 nm, tiled 2×2 to acquire the entire well. For *S. mansoni* adult motility, wells were acquired for 60 frames at 2x with transmitted light and 2x binning.

Raw images were exported with MetaXpress v6 and stored on the UW-Madison Research Drive in an uncompressed state. When analyzed using a distributed computing system, images were transferred to the UW-Madison Center for High-Throughput Computing (CHTC) submit servers. Jobs were submitted and managed with HTCondor [16]. HTCondor submit scripts are publicly available at https://github.com/zamanianlab/chtc-submit/tree/main/imgproc.

### Worm classification model building

Straightened worm images from 6 different experiments were manually classified as Single worm, Partial worm, Multiple worms, or Debris (8535 objects in total). Each object was associated with 25 area/shape features and 15 intensity features output by CellProfiler, and the features and training data were used to build several classification models using a variety of approaches, including the boosted frameworks XGBoost [17,18] and LightGBM [19]. These were implemented using the tidymodels package in the R statistical software [20]. Highly correlated predictors and predictors with near-zero variance were removed. Care was taken to attempt to balance the dataset by worm classification and the size of single worms, and lowly represented classes were upsampled with the SMOTE method [21]. Data was split into training/testing sets, and the training results were evaluated using 10-fold cross validation. Model hyperparameters were tuned using grid searches. Tuned models were fit to the cross-fold testing set and evaluated using the area under the receiver-operator curve (ROC AUC).

## Results

### wrmXpress is user-friendly, modular, and extensible

wrmXpress is a unified framework for analyzing worm imaging data. It comes packaged with a variety of phenotyping modules matched to specific experimental setups, including motility and viability for parasites and *C. elegans*, and feeding rate and development in *C. elegans*. The package is easily extensible using open-source Python libraries or new CellProfiler pipelines.

The package combines code, user-generated job parameters, and input data/metadata (Fig 1), each of which is passed to a Docker container that runs the pipeline. It is implemented with a single command (e.g., `python wrmXpress/wrapper.py params.yml {plate_dir}`), and modules are configured and initialized with the YAML parameters file that designates the species, worm stage, and the modules to be run (Fig 2A). A user can choose to analyze the entire 96-well plate or a selected subset of wells. The wrapper.py script integrates these selections and runs the proper modules and commands (Fig 2B). Output data is written to a directory with raw output, a tidied output that joins well-based experimental metadata, and thumbnail images to assist with quality control and error diagnosis (Fig 2C).

**Figure 1.**
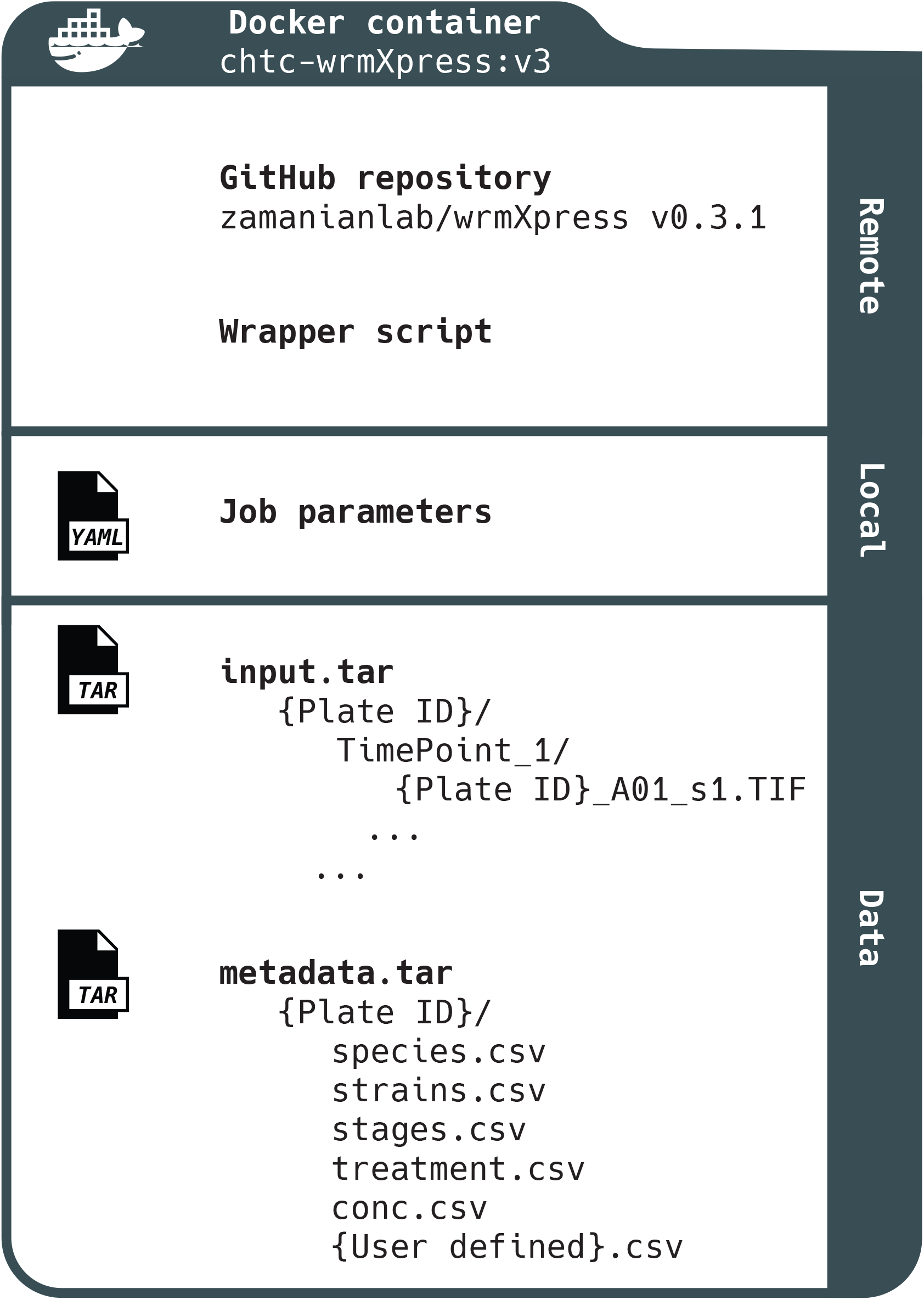
Schematic of wrmXpress. wrmXpress consists of code that is held in a public GitHub repository (including the master wrapper script), job parameters that are edited locally, and external data and metadata. The structure of input data and metadata requires specific formats in order for wrmXpress to complete without error.

**Figure 2.**
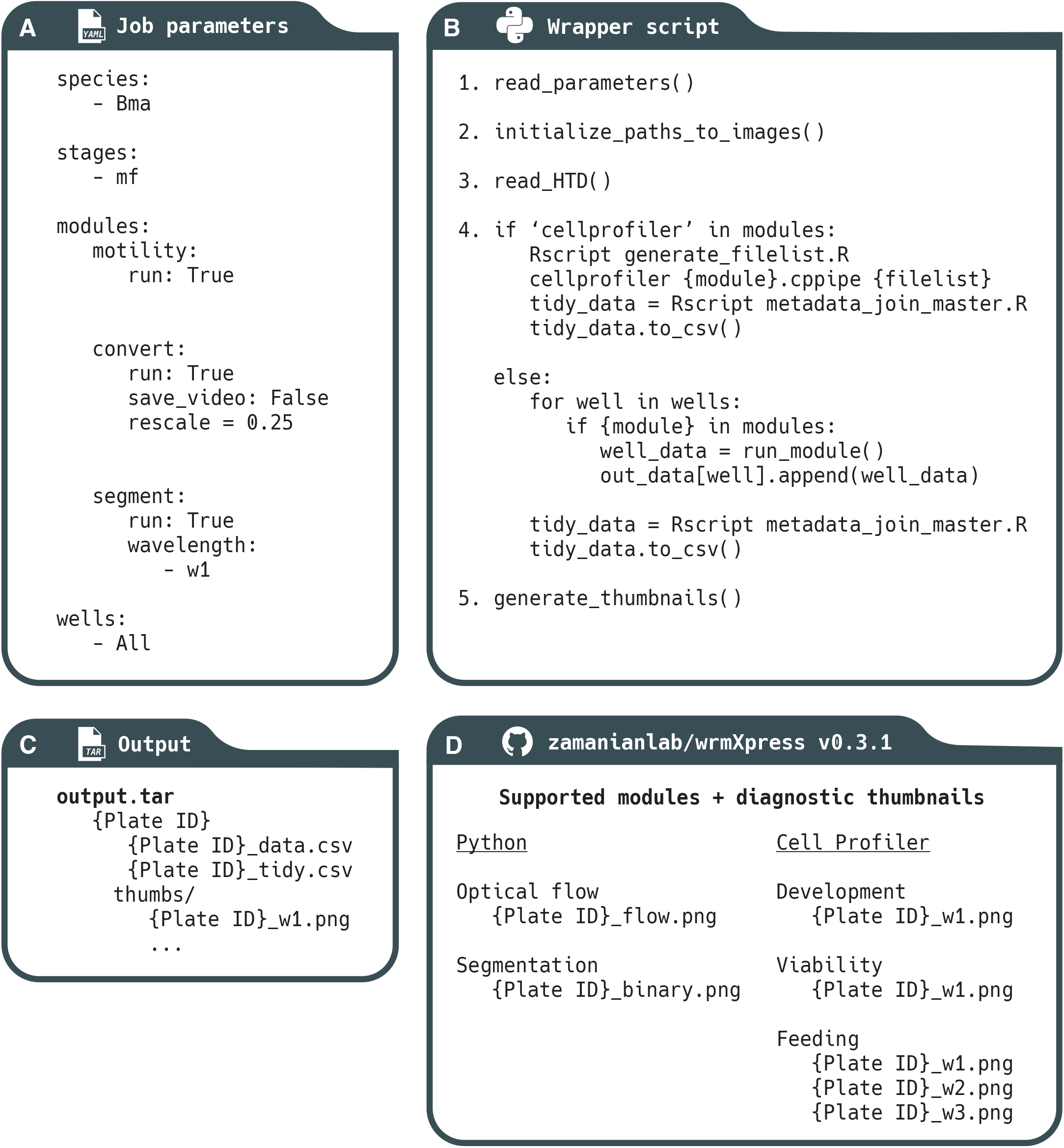
Constituents of the wrmXpress workflow. (A) Jobs are parameterized with a user-generated YAML file, which includes species and stage information, and allows for the selection of Python or CellProfiler modules. (B) The wrapper scripts control the implementation of wrmXpress. (C) Output data includes raw data, raw data with joined metadata, and diagnostic thumbnail images. (D) wrmXpress comes packaged with 5 distinct analytical modules.

### wrmXpress usage

wrmXpress is designed such that each module outputs a single phenotypic value per well, or multiple values per well or object if using a CellProfiler pipeline. For instance, for a motility experiment that could use worm area as a normalization coefficient, both the motility and segmentation modules can be selected, which will calculate the raw optical flow and total worm area per well. Each value is then concatenated to a final output file that includes metadata and per-module measurements.

At the start of a wrmXpress run, the user-generated parameters provided by the YAML are read and organized (Fig 2B, step 1). Paths to relevant image and metadata files are populated, and modules are selected (Fig 2B, step 2). It is during this stage that wells of interest can be selected in order to reduce runtime in case of contamination, empty wells, or during testing. Since not all modules are compatible (for instance, some require multiple time points and some require multiple wavelengths), some light checking of parameters and input data is performed in order to avoid module clashes and to ensure a correct pairing between modules and input data. Finally, the plate’s HTD file, a machine-generated configuration file that reports imager settings, is parsed (Fig 2B, step 3). These imager configurations are used in some downstream modules, like stitching of tiled images.

Once paths and parameters are organized, the wrapper script loops through the selected wells and iteratively calls the functions for each selected module (Fig 2B, step 4). For Cell Profiler pipelines, an R script utilizes the populated file paths to automatically generate the CSV that is used by CellProfiler’s LoadData module. CellProfiler is then called in headless mode. Each pipeline must also include the ExportToSpreadsheet module, which collects the well and/or object-based data and writes it to a CSV. Finally, another R script joins user-provided experimental metadata to the output CSV to create a final tidy data file.

For bespoke Python modules, less preparation is required. As the wrapper iterates through wells, each module is called independently of other modules. After completion of a well, the module will return a single phenotypic value, which is added to a dictionary of values that is dynamically updated. After iterating through all selected wells, the dictionary is written to a CSV, and the data is tidied as above.

After Cell Profiler pipelines or Python modules are finished, diagnostic thumbnails are generated and formatted in a 12×8 array. By default, a thumbnail will be created for each included wavelength, and specific modules generate relevant diagnostics to help evaluate module performance.

### Analytical modules for worm motility, area, development, viability and feeding

wrmXpress comes packaged with five individual modules that enable a wide range of out-of-the-box functionalities (Fig 2D). Motility measurements are implemented using a dense measure of optical flow (Farneback’s method [22]). Dense flow for worm motility has been used elsewhere [10], and offers a richer output than previous optical flow-based implementations that prioritized a sparse feature set (the Lucas-Kanade method [23]). Focusing on a sparse feature set enabled real-time tracking of worms [3,4], but given that real-time tracking is not a priority in high-throughput approaches, wrmXpress opts for the more data-rich option. A unitless measure of motility is calculated by summing the magnitude of the flow vectors across n-1 frames, and then summing the sum across all pixels. Thus, flow is a function of video length as well as each frame’s height and width. This algorithm has been tested for *Brugia* spp. microfilariae, *C. elegans* L1s and adults, and *S. mansoni* adult males and females (Fig 3A-C). A “flow cloud” diagnostic thumbnail is created for each well, providing an image representation of the motility over the entire course of the video. When multiple worms are included per well, this diagnostic can also alert investigators to plate effects or heterogeneity between wells.

**Figure 3.**
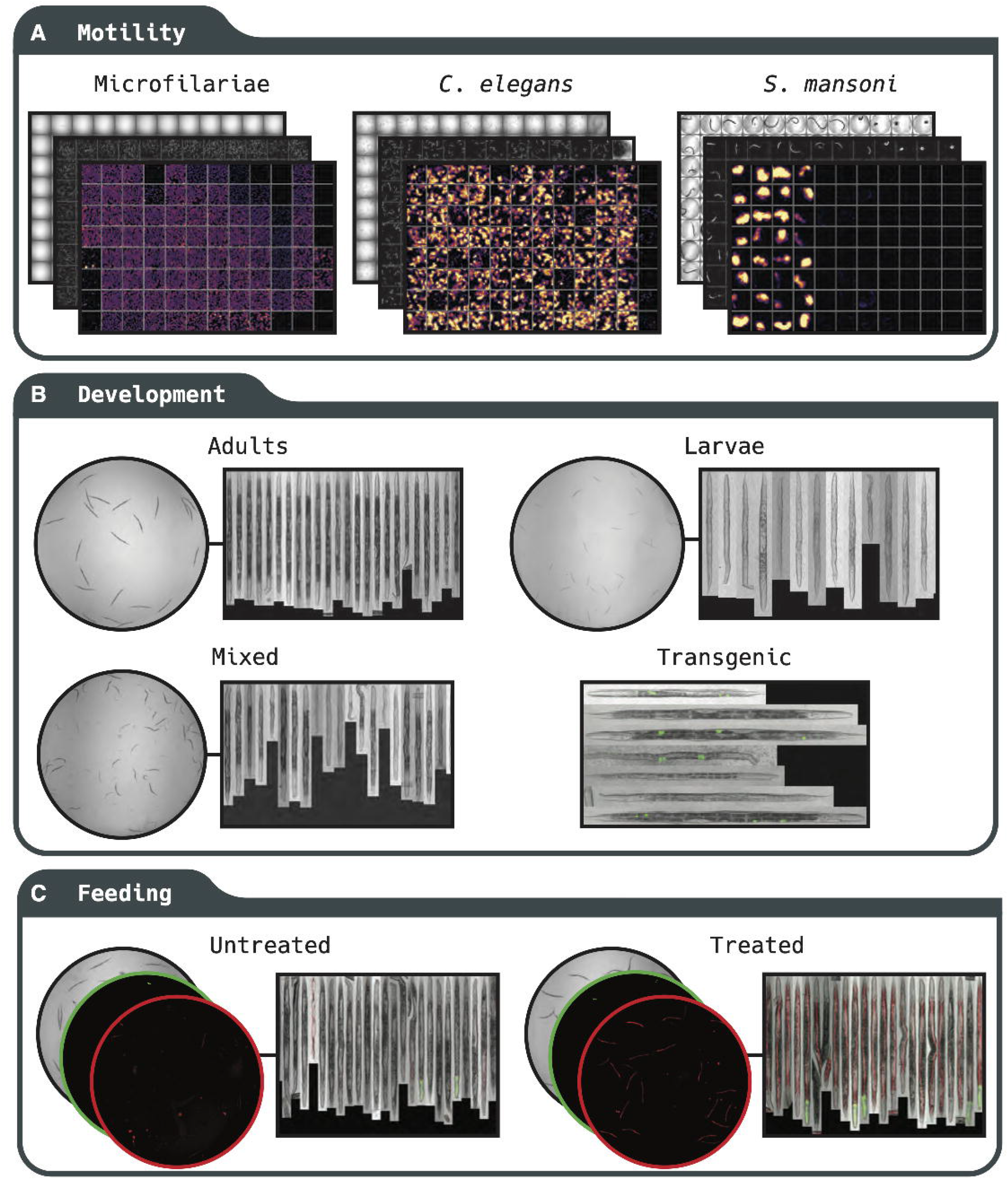
Examples of phenotypes that can be analyzed with wrmXpress. (A) Motility of *B. malayi* microfilariae, *C. elegans* adults, and *S. mansoni* adults. Diagnostic images include a single frame of transmitted light, binary segmented worms, and a flow cloud. (B) Development of *C. elegans*, which is amenable for staged adults and larvae, as well as wells with mixed populations. Classification of transgenic worms (*unc-122p::GFP*) is also implemented. (C) Quantification of *C. elegans* feeding using fluorescent dies, which can be measured in the worm intestine.

Motility measurements on wells with multiple worms can be normalized by dividing the motility value by the worm area, which is calculated by the segmentation module. We have found that a simple algorithm incorporating Sobel edge detection, Gaussian blur, and Otsu’s thresholding method performs well for a variety of vermiform objects (including all nematodes so far tested) [24,25]. For larger worms that are less optically translucent (e.g., *S. mansoni* adults), we implement Gaussian blur followed by a simple percentile threshold. The percentile and σ for the Gaussian kernel may need to be adjusted in accordance with varying illumination parameters, but the defaults have been robust in our hands (1.5% and σ = 1.5). For *S. mansoni* adult females or male/female pairs, which eject a variety of debris in culture, an object size filter has been implemented. The final binary segmented image is also written out as a diagnostic thumbnail (Fig 3).

We have used a similar segmentation algorithm for the quantification of female *Brugia* spp. fecundity. Though a fecundity module has not technically been integrated into wrmXpress, the segmentation module can be used for a variety of end user needs.

Integration of CellProfiler pipelines further extends the capabilities of wrmXpress, which comes with pre-built pipelines for analyzing *C. elegans* development and feeding, each of which can be used on mixed populations of transgenic worms with a fluorescent marker (Fig 3B-C). These pipelines take advantage of the WormToolbox plugin, which incorporates user-generated worm models to segment and untangle individual worms [11]. For the wrmXpress development module, a number of innovations were necessary to prepare the pipeline for identifying and retaining worms that greatly varied in size, as drug treatment of synchronized worms can lead to mixed populations of worms in a single well (Fig 3B). Relaxation of the segmentation algorithm predictably led to the inclusion of more debris as objects, so we trained a post-processing classification model that used object shape and intensity features as predictors to classify untangled worms as a single worm, partial worm, multiple worms, or debris. We tuned, trained, and evaluated a variety of machine learning models and selected a gradient boosted tree due to its performance and speed of classification on experimental data (Fig 4A). The trained model used the minimum y (length) of the bounding box, the minimum and standard deviation of the intensity of the edge pixels, and the solidity (the ratio of the contour area to its convex hull area), among others, as the most important variables (Fig 4B). The model achieved an area under the receiver-operator characteristic curve (AUC ROC) of 0.827 and a sensitivity of 0.609 (S1 File). When fit to annotated holdout data and retaining only those predicted to be single worms, the model removed 93% of the debris, 89% of multiple worms, 93% of partial worms, and only 18% of single worms, substantially enriching for objects of interest. In our hands, using a single, less stringent worm model in CellProfiler followed by post-processing filtration decreased runtime and ensured that smaller worms were captured. Both models (the worm model used in CellProfiler and the classification model used in post-processing) should be trained on user-generated data. Instructions for training the former are available on the CellProfiler documentation website, and we have included example pipelines for selecting worms and training the model in the GitHub repository.

**Figure 4.**
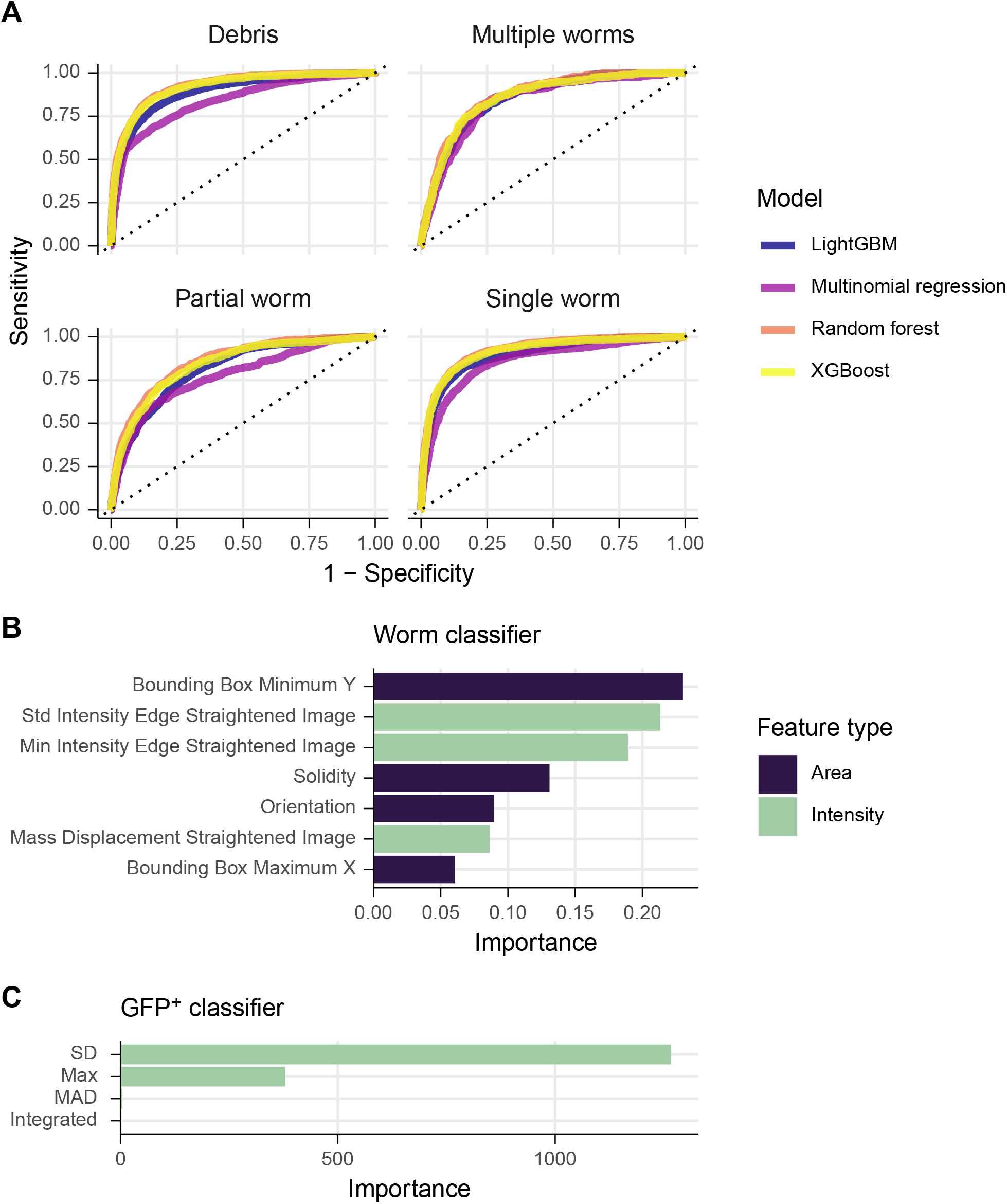
Statistical models used to classify and filter straightened *C. elegans*. (A) Evaluation of models classifying segmented and straightened “worms” as debris, multiple worms, a partial worm, or a single worm. (B) Variable importance plot for the tuned XGBoost model from (A). Green bars indicate intensity features, purple indicate area/shape features.l (C) Variable importance plot for a tuned random forest to classify GFP+/-worms (*unc-122p::GFP*).

For transgenic worms with fluorescent markers, we also chose to filter during post-processing rather than implementing a filter in the CellProfiler pipeline. This allows for labeling each worm as +/- in the final tidied data, providing a convenient within-well control population (transgenic strains generated with extrachromosomal arrays contain a mix of transgene^+^ and transgene^-^ worms). We trained a simple random forest on annotated worms that were labeled with *unc-122p::GFP*, which is fluorescent in only a handful of cells (Fig 4B). This classifier achieved 100% accuracy, and the most important variable in the model was the standard deviation of the fluorescence intensity (Fig 5C). Internally, we also use this model for classifying pharynx-labeled (*myo-2p::GFP*) transgenic worms.

Finally, we include a CellProfiler pipeline for the measurement of staining by a viability dye (CellTox), which we have used with both microfilariae and adult *C. elegans.* This pipeline uses similar principles to the segmentation module but is optimized for fluorescent images. The pipeline will segment stained areas and output a measure of total fluorescence.

### wrmXpress is readily extensible

wrmXpress can be extended by developing new, isolated modules (e.g., Python scripts) that take the images from a single well, perform transformations/calculations on them, and output a single value. For instance, one can easily imagine a Python module that counts segmented objects in a well. The Python script can be written, added to the `modules`/ directory, added to the if/else loop in the wrapper script, and added as an option in the YAML configuration template. The module will be run independently, enabling safe, backwards-compatible engineering of new modules.

Likewise, new CellProfiler pipelines can also be easily implemented. In this case, a pipeline is developed in the CellProfiler GUI, exported as a .cppipe file, added to the `cp_pipelines/pipelines/` directory, and added as an option in the YAML configuration template. As an additional step, a user must also add an R script that parses the input file names and generates the CSV file that is read by the LoadData module in CellProfiler.

wrmXpress does not have a GUI and therefore can only be extended by R and Python developers. However, we have taken great pains to make the addition of Python modules or CellProfiler pipelines simple and barrier-free. Additionally, we have found that researchers without programming experience can develop pipelines using the CellProfiler GUI, which can then be integrated into the wrmXpress framework by novice developers. Lab specific documentation for extending wrmXpress can be found at http://www.zamanianlab.org/ZamanianLabDocs/pipelines_wrmxpress/, which may be instructive.

## Conclusions and future developments

We view wrmXpress as a part of the next-generation of parasitic worm phenotyping toolkits, building upon important advances made by WormAssay/Worminator[3,4] and the WormToolbox[11] and enabled by high-content imaging. The software contains a variety of analytical modules that are optimized for experiments with worms, and new modules are being developed. For instance, we are actively experimenting with methods for quantifying adult worm fecundity by segmenting adults and progeny and using machine learning classifiers to count distinct classes of objects. We believe this approach will be easily adapted for *C. elegans* and *S. mansoni* and could make use of new culture media that enable *in vitro* fecundity [26].

Future developments in high-content phenotyping of worms likely include the utilization of deep learning frameworks for a variety of phenotypic endpoints. We have observed that drug treatment of worms can cause diverse, often ephemeral, motile behaviors that can be identified by eye, but as of yet cannot be classified computationally [10]. We have additionally observed that drug-induced worm death can result in one of a number of different worm postures, which we believe is related to drug mechanisms of action, in the same way that drug MoA can be parsed by classifying behavioral fingerprints in *C. elegans* [27]. Deep learning is well suited for each of these tasks, and the structure of wrmXpress is such that deep learning modules can easily be added. Indeed, these extensions are actively being developed.

Due to limitations in running CellProfiler in headless mode, wrmXpress cannot currently be run in parallel (i.e., analyzing individual wells by separate processors). However, high-throughput screens often generate dozens of plates per day, and wrmXpress is readily capable of analyzing plates in parallel by submitting separate jobs for each plate (or running separate commands on a local machine). Indeed, this is our current implementation with HTCondor [16]. However, future developments of wrmXpress could allow for well-based parallelization, either by changes to the handling and organization of input data, or by making use of Python’s multiple libraries for parallelization. Regardless, in our hands the analysis of a full 96-well plate takes less than 3 hours using relatively modest hardware specifications (4 CPUs, 20 GB RAM).

wrmXpress will work out-of-the-box for all datasets generated with an ImageXpress (Molecular Devices), which is a popular platform for worm labs [13,15,28,29]. For other endpoints, an HTD file must be provided, and the image data must be structured as in Fig 1. However, the design of the pipelines is such that adding support for other platforms will be straightforward.

wrmXpress v1.0.0 can be downloaded from its public GitHub repository (https://github.com/zamanianlab/wrmXpress), and the Docker container that includes all dependencies is also available (https://hub.docker.com/r/zamanianlab/chtc-wrmxpress). Documentation can be found at the GitHub repository, and additional developer information can be found at http://www.zamanianlab.org/ZamanianLabDocs/pipelines_wrmxpress/.

## Acknowledgements

This research was performed using the compute resources and assistance of the UW-Madison Center For High Throughput Computing (CHTC) in the Department of Computer Sciences. The CHTC is supported by UW-Madison, the Advanced Computing Initiative, the Wisconsin Alumni Research Foundation, the Wisconsin Institutes for Discovery, and the National Science Foundation, and is an active member of the OSG Consortium, which is supported by the National Science Foundation and the U.S. Department of Energy’s Office of Science.

## Funding

This work was supported by National Institutes of Health NIAID grant R01 AI151171 to MZ and R21 AI153545 to MZ and JDC. NJW was supported by NIH Ruth Kirschstein NRSA fellowship F32 AI152347.

## Supporting information captions

S1 File. Building and evaluating models for classifying straightened worm objects.

## Notes

### Competing Interest Statement

The authors have declared no competing interest.

## References

1. Zamanian M, Chan JD. High-content approaches to anthelmintic drug screening. Trends Parasitol. 2021;37: 780–789.

2. Herath HMPD, Taki AC, Rostami A, Jabbar A, Keiser J, Geary TG, et al. Whole-organism phenotypic screening methods used in early-phase anthelmintic drug discovery. Biotechnol Adv. 2022;57: 107937.

3. Marcellino C, Gut J, Lim KC, Singh R, McKerrow J, Sakanari J. WormAssay: a novel computer application for whole-plate motion-based screening of macroscopic parasites. PLoS Negl Trop Dis. 2012;6: e1494.

4. Storey B, Marcellino C, Miller M, Maclean M, Mostafa E, Howell S, et al. Utilization of computer processed high definition video imaging for measuring motility of microscopic nematode stages on a quantitative scale: “The Worminator.” Int J Parasitol Drugs Drug Resist. 2014;4: 233–243.

5. Partridge FA, Brown AE, Buckingham SD, Willis NJ, Wynne GM, Forman R, et al. An automated high-throughput system for phenotypic screening of chemical libraries on C. elegans and parasitic nematodes. Int J Parasitol Drugs Drug Resist. 2018;8: 8–21.

6. Preston S, Jabbar A, Nowell C, Joachim A, Ruttkowski B, Baell J, et al. Low cost whole-organism screening of compounds for anthelmintic activity. Int J Parasitol. 2015;45: 333–343.

7. Ritler D, Rufener R, Sager H, Bouvier J, Hemphill A, Lundström-Stadelmann B. Development of a movement-based in vitro screening assay for the identification of new anti-cestodal compounds. PLoS Negl Trop Dis. 2017;11: e0005618.

8. Weeks JC, Roberts WM, Leasure C, Suzuki BM, Robinson KJ, Currey H, et al. Sertraline, Paroxetine, and Chlorpromazine Are Rapidly Acting Anthelmintic Drugs Capable of Clinical Repurposing. Sci Rep. 2018;8: 1–17.

9. Tyagi R, Bulman CA, Cho-Ngwa F, Fischer C, Marcellino C, Arkin MR, et al. An Integrated Approach to Identify New Anti-Filarial Leads to Treat River Blindness, a Neglected Tropical Disease. Pathogens. 2021;10. doi:10.3390/pathogens10010071

10. Wheeler NJ, Heimark ZW, Airs PM, Mann A, Bartholomay LC, Zamanian M. Genetic and functional diversification of chemosensory pathway receptors in mosquito-borne filarial nematodes. PLoS Biol. 2020;18: e3000723.

11. Wählby C, Kamentsky L, Liu ZH, Riklin-Raviv T, Conery AL, O’Rourke EJ, et al. An image analysis toolbox for high-throughput C. elegans assays. Nat Methods. 2012;9: 714–716.

12. McQuin C, Goodman A, Chernyshev V, Kamentsky L, Cimini BA, Karhohs KW, et al. CellProfiler 3.0: Next-generation image processing for biology. PLoS Biol. 2018;16: e2005970.

13. Paveley RA, Mansour NR, Hallyburton I, Bleicher LS, Benn AE, Mikic I, et al. Whole organism high-content screening by label-free, image-based Bayesian classification for parasitic diseases. PLoS Negl Trop Dis. 2012;6: e1762.

14. Chen S, Suzuki BM, Dohrmann J, Singh R, Arkin MR, Caffrey CR. A multi-dimensional, time-lapse, high content screening platform applied to schistosomiasis drug discovery. Commun Biol. 2020;3: 747.

15. Nyaanga J, Crombie TA, Widmayer SJ, Andersen EC. easyXpress: An R package to analyze and visualize high-throughput C. elegans microscopy data generated using CellProfiler. PLoS One. 2021;16: e0252000.

16. Thain D, Tannenbaum T, Livny M. Distributed computing in practice: the Condor experience. Concurr Comput. 2005;17: 323–356.

17. Chen T, Guestrin C. XGBoost: A Scalable Tree Boosting System. Proceedings of the 22nd ACM SIGKDD International Conference on Knowledge Discovery and Data Mining. New York, NY, USA: Association for Computing Machinery; 2016. pp. 785–794.

18. Chen T, He T, Benesty M, Khotilovich V, Tang Y, Cho H, et al. Xgboost: extreme gradient boosting. R package version 0 4-2. 2015;1: 1–4.

19. Ke G, Meng Q, Finley T, Wang T, Chen W, Ma W, et al. LightGBM: A highly efficient gradient boosting decision tree. Adv Neural Inf Process Syst. 2017;30. Available: https://proceedings.neurips.cc/paper/2017/hash/6449f44a102fde848669bdd9eb6b76fa-Abstract.html

20. Kuhn M, Wickham H. Tidymodels: a collection of packages for modeling and machine learning using tidyverse principles. Boston, MA, USA [(accessed on 10 December 2020)]. 2020.

21. Chawla NV, Bowyer KW, Hall LO, Kegelmeyer WP. SMOTE: Synthetic Minority Over-sampling Technique. J Artif Intell Res. 2002;16: 321–357.

22. Farnebäck G. Two-Frame Motion Estimation Based on Polynomial Expansion. Image Analysis. Springer Berlin Heidelberg; 2003. pp. 363–370.

23. Lucas BD, Kanade T. An iterative image registration technique with an application to stereo vision. Proceedings of the 7th international joint conference on Artificial intelligence - Volume 2. San Francisco, CA, USA: Morgan Kaufmann Publishers Inc.; 1981. pp. 674–679.

24. Otsu N. A Threshold Selection Method from Gray-Level Histograms. IEEE Trans Syst Man Cybern. 1979;9: 62–66.

25. Kanopoulos N, Vasanthavada N, Baker RL. Design of an image edge detection filter using the Sobel operator. IEEE J Solid-State Circuits. 1988;23: 358–367.

26. Wang J, Chen R, Collins JJ 3rd. Systematically improved in vitro culture conditions reveal new insights into the reproductive biology of the human parasite Schistosoma mansoni. PLoS Biol. 2019;17: e3000254.

27. McDermott-Rouse A, Minga E, Barlow I, Feriani L, Harlow PH, Flemming AJ, et al. Behavioral fingerprints predict insecticide and anthelmintic mode of action. Mol Syst Biol. 2021;17: e10267.

28. Edwards J, Brown M, Peak E, Bartholomew B, Nash RJ, Hoffmann KF. The diterpenoid 7-keto-sempervirol, derived from Lycium chinense, displays anthelmintic activity against both Schistosoma mansoni and Fasciola hepatica. PLoS Negl Trop Dis. 2015;9: e0003604.

29. Giuliani S, Silva AC, Borba JVVB, Ramos PIP, Paveley RA, Muratov EN, et al. Computationally-guided drug repurposing enables the discovery of kinase targets and inhibitors as new schistosomicidal agents. PLoS Comput Biol. 2018;14: e1006515.

