## Supplementary material for "wrmXpress: A modular package for high-throughput image analysis of parasitic and free-living worms": S1 File

### Models for filtering Worm Toolbox output

## 2022-05-20

This document assesses the ability of several models to classify StraightenedWorm data. Manual annotations are used to train, validate, and select the best model and hyperparameters

First load the data:

```
library(tidymodels)
library(tidyverse)
library(treesnip)
library(here)

path <- list.files(path = here(),
                   pattern = '.*(EJG|NJW).*_tidy.csv',
                   recursive = TRUE)

annotations <- read_csv(here(path)) %>%
  janitor::clean_names() %>%
  filter(!is.na(category)) %>%
  select(category, contains(c('area', 'intensity'))) %>%
  mutate(category = case_when(
    category == 'Y' ~ 'Single worm',
    category == 'D' ~ 'Debris',
    category == 'P' ~ 'Partial worm',
    category == 'M' ~ 'Multiple worms',
    FALSE ~ 'Other'
  )) %>%
  filter(category != 'Other') %>%
  mutate(category = as.factor(category))

glimpse(annotations)
```

```
## Rows: 8,535
## Columns: 41
## $ category                                            <fct> Debris, Single …
## $ area_shape_area                                     <dbl> 712, 2029, 1300…
## $ area_shape_bounding_box_area                        <dbl> 798, 2205, 1428…
## $ area_shape_bounding_box_maximum_x                   <dbl> 21, 42, 63, 84,…
## $ area_shape_bounding_box_maximum_y                   <dbl> 40, 109, 72, 85…
## $ area_shape_bounding_box_minimum_x                   <dbl> 0, 21, 42, 63, …
## $ area_shape_bounding_box_minimum_y                   <dbl> 2, 4, 4, 4, 4, …
## $ area_shape_center_x                                 <dbl> 9.707865, 31.01…
## $ area_shape_center_y                                 <dbl> 20.69803, 55.92…
## $ area_shape_compactness                              <dbl> 1.169305, 2.108…
## $ area_shape_convex_area                              <dbl> 723, 2053, 1318…
## $ area_shape_eccentricity                             <dbl> 0.8243468, 0.97…
## $ area_shape_equivalent_diameter                      <dbl> 30.10891, 50.82…
## $ area_shape_euler_number                             <dbl> 1, 1, 1, 1, 1, …
## $ area_shape_extent                                   <dbl> 0.8922306, 0.92…
## $ area_shape_form_factor                              <dbl> 0.8552088, 0.47…
## $ area_shape_major_axis_length                        <dbl> 40.24201, 112.6…
## $ area_shape_max_feret_diameter                       <dbl> 37.85499, 104.0…
## $ area_shape_maximum_radius                           <dbl> 11, 11, 11, 11,…
## $ area_shape_mean_radius                              <dbl> 4.774433, 5.352…
## $ area_shape_median_radius                            <dbl> 4.123106, 5.000…
## $ area_shape_min_feret_diameter                       <dbl> 20, 20, 20, 20,…
## $ area_shape_minor_axis_length                        <dbl> 22.78041, 23.35…
## $ area_shape_orientation                              <dbl> 3.572102e+00, -…
## $ area_shape_perimeter                                <dbl> 102.28427, 231.…
## $ area_shape_solidity                                 <dbl> 0.9847856, 0.98…
## $ intensity_integrated_intensity_edge_straightened_image <dbl> 2.317286, 8.323…
## $ intensity_integrated_intensity_straightened_image     <dbl> 23.44981, 73.16…
## $ intensity_lower_quartile_intensity_straightened_image  <dbl> 0.03216602, 0.0…
## $ intensity_mad_intensity_straightened_image           <dbl> 0.001220722, 0.…
## $ intensity_mass_displacement_straightened_image       <dbl> 0.14369717, 0.3…
## $ intensity_max_intensity_edge_straightened_image      <dbl> 0.03718623, 0.0…
## $ intensity_max_intensity_straightened_image           <dbl> 0.03718623, 0.0…
## $ intensity_mean_intensity_edge_straightened_image     <dbl> 0.03458635, 0.0…
## $ intensity_mean_intensity_straightened_image          <dbl> 0.03293512, 0.0…
## $ intensity_median_intensity_straightened_image        <dbl> 0.03361563, 0.0…
## $ intensity_min_intensity_edge_straightened_image      <dbl> 0.025360495, 0.…
## $ intensity_min_intensity_straightened_image           <dbl> 0.023702238, 0.…
## $ intensity_std_intensity_edge_straightened_image      <dbl> 0.002046304, 0.…
## $ intensity_std_intensity_straightened_image           <dbl> 0.002676937, 0.…
## $ intensity_upper_quartile_intensity_straightened_image  <dbl> 0.03474479, 0.0…
```

### Explore data

The Worm Toolbox in Cell Profiler can export a variety of features, some of which may be useful in classification.

```r
library(ggbeeswarm)

annotations %>%
  pivot_longer(-category, names_to = 'measurement', values_to = 'value') %>%
  ggplot() +
  geom_quasirandom(aes(x = category, y = value, color = category)) +
  facet_wrap(facets = vars(measurement), scales = 'free_y') +
  theme_minimal() +
  NULL
```

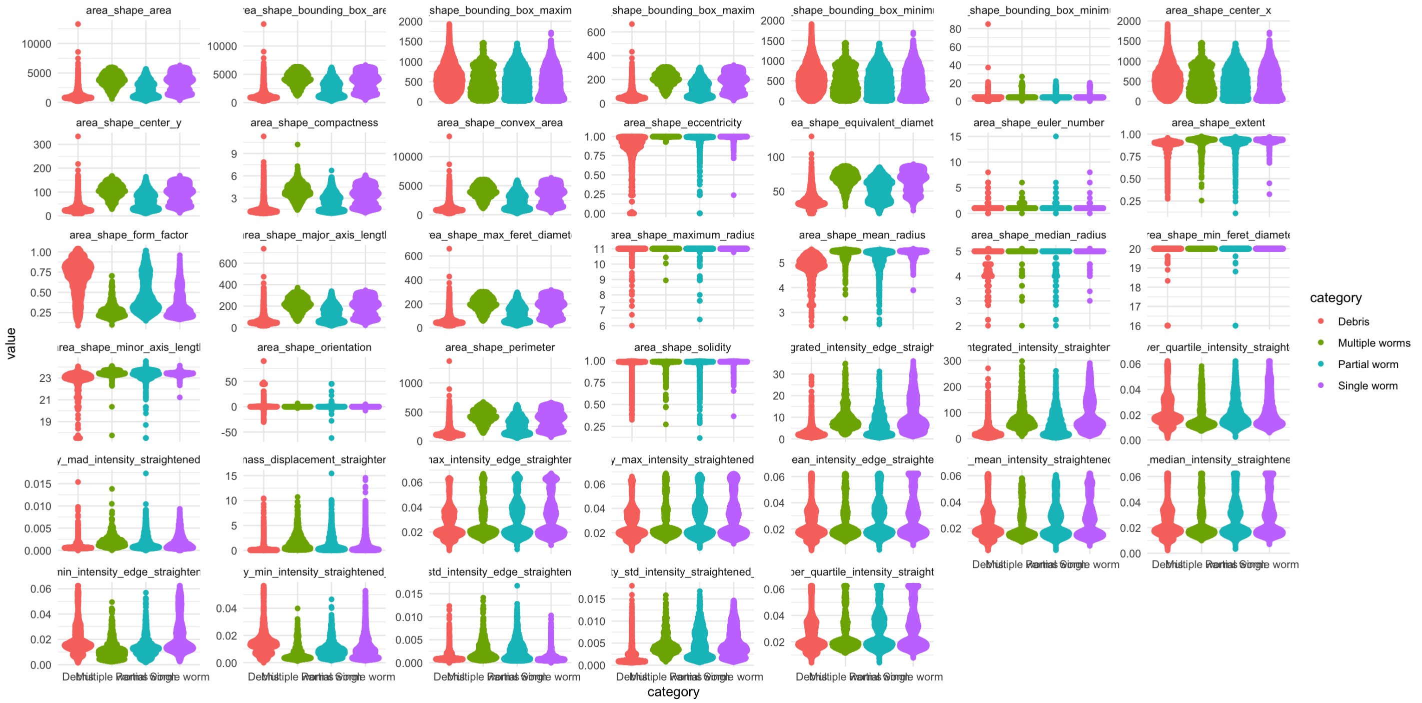


### Build models

First create training (with cross-fold validation) and test data sets

```r
model_data <- annotations %>%
  mutate(category = factor(category))

set.seed(123)
# data_boot <- bootstraps(model_data, times = 2) # only 2 bootstraps for testing
data_split <- initial_split(model_data,
                            strata = category)
train_data <- training(data_split)
test_data <- testing(data_split)

set.seed(234)
folds <- vfold_cv(train_data,
                  v = 10,
                  strata = category)
```

Specify the models:

```r
library(doParallel)
cores <- parallel::detectCores(logical = FALSE)
registerDoParallel(cores = 4)

multinom_reg_glmnet_spec <-
  multinom_reg(penalty = tune(), mixture = tune()) %>%
  set_engine('glmnet')

rand_forest_ranger_spec <-
  rand_forest(mtry = tune(), min_n = tune()) %>%
  set_engine('ranger', num.threads = cores) %>%
  set_mode('classification')

boost_tree_xgboost_spec <-
  boost_tree(tree_depth = tune(), learn_rate = tune(),
             min_n = tune(), loss_reduction = tune(), mtry = tune(),
             sample_size = tune(), stop_iter = tune()) %>%
  set_engine('xgboost') %>%
  set_mode('classification')

light_gbm_spec <-
  boost_tree(trees = 1000,  min_n = tune(),
             tree_depth = tune(), learn_rate = tune(),
             loss_reduction = tune()) %>%
  set_engine('lightgbm') %>%
  set_mode('classification')
```

Build the recipe and workflow:

```r
library(themis)

recipe <-
  recipe(category ~ ., data = model_data) %>%
  step_nzv(all_predictors()) %>%
  step_normalize(all_predictors()) %>%
  step_corr(all_numeric_predictors(), threshold = .5) %>%
  step_smote(category)

prep <- prep(recipe)
juice <- juice(prep)

prep
```

```
## Recipe
##
## Inputs:
##
##       role #variables
##    outcome          1
##  predictor         40
##
## Training data contained 8535 data points and no missing data.
##
## Operations:
##
## Sparse, unbalanced variable filter removed area_shape_euler_number, area_shape_max...
[trained]
## Centering and scaling for area_shape_area, area_shape_bounding_box_area, ... [traine
d]
## Correlation filter on area_shape_bounding_box_maximum_y, area_s... [trained]
## SMOTE based on category [trained]
```

```
glimpse(juice)
```

```
## Rows: 14,476
## Columns: 8
## $ area_shape_bounding_box_maximum_x               <dbl> -1.539705342, -1.47806…
## $ area_shape_bounding_box_minimum_y               <dbl> -0.79148943, -0.042184…
## $ area_shape_orientation                          <dbl> 0.61686797, -0.1169251…
## $ area_shape_solidity                             <dbl> 0.24346173, 0.29945416…
## $ intensity_mass_displacement_straightened_image  <dbl> -0.61610872, -0.498939…
## $ intensity_min_intensity_edge_straightened_image <dbl> 0.52530035, 0.32174241…
## $ intensity_std_intensity_edge_straightened_image <dbl> 0.278401463, 0.0568308…
## $ category                                        <fct> Debris, Single worm, S…
```

```
recipe2 <- recipe
recipe2$steps[[3]] <- update(recipe2$steps[[3]], skip = TRUE)

mn_workflow <-
  workflow() %>%
  add_model(multinom_reg_glmnet_spec) %>%
  add_recipe(recipe)

rf_workflow <-
  workflow() %>%
  add_model(rand_forest_ranger_spec) %>%
  add_recipe(recipe2)

xg_workflow <-
  workflow() %>%
  add_model(boost_tree_xgboost_spec) %>%
  add_recipe(recipe)

lgbm_workflow <-
  workflow() %>%
  add_model(light_gbm_spec) %>%
  add_recipe(recipe)
```

### Tune the models

Uncomment the pipes that include `tune_grid()` to actually perform the turning. For simplicity here, tuning was done separately and the resulting data is read as an RDS file for evaluation.

#### Multinomial regression

```r
mn_grid <- grid_regular(mixture(),
                        penalty())

# mn_tune <-
#   mn_workflow %>%
#   tune_grid(
#     resamples = folds,
#     grid = mn_grid,
#     control = control_grid(save_pred = TRUE,
#                            verbose = TRUE),
#     metrics = metric_set(roc_auc, sens))
#
# write_rds(mn_tune, here('code', 'rds', 'mn_tune.rds'))

mn_tune <- read_rds(here('code', 'rds', 'mn_tune.rds'))

# extract the best model
best_mn <- mn_tune %>%
  select_best("roc_auc")

# print metrics
(mn_metrics <- mn_tune %>%
    collect_metrics() %>%
    semi_join(best_mn) %>%
    select(.metric:.config) %>%
    mutate(model = 'Multinomial regression'))
```

```
## # A tibble: 2 × 7
##   .metric .estimator  mean     n std_err .config              model
##   <chr>   <chr>      <dbl> <int>   <dbl> <chr>                <chr>
## 1 roc_auc hand_till  0.802    10 0.00386 Preprocessor1_Model7 Multinomial regre…
## 2 sens    macro      0.570    10 0.00746 Preprocessor1_Model7 Multinomial regre…
```

```r
# finalize the wf with the best model
mn_workflow <-
  mn_workflow %>%
  finalize_workflow(best_mn)

# generate predictions on the hold-out test data
mn_auc <-
  mn_tune %>%
  collect_predictions(parameters = best_mn) %>%
  roc_curve(.pred_Debris:`.pred_Single worm`, truth = category) %>%
  mutate(model = "Multinomial regression")

mn_auc %>%
  autoplot()
```

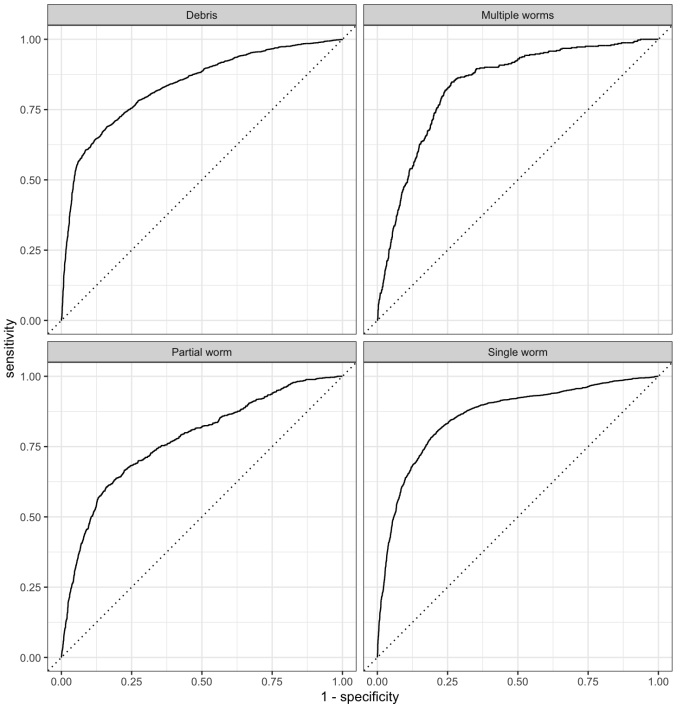


### Random forest

```
rf_grid <- grid_regular(finalize(mtry(), model_data),
                        min_n())

# rf_tune <-
#   rf_workflow %>%
#   tune_grid(
#     resamples = folds,
#     grid = rf_grid,
#     control = control_grid(save_pred = TRUE,
#                            verbose = TRUE),
#     metrics = metric_set(roc_auc, sens))
#
# write_rds(rf_tune, here('code', 'rds', 'rf_tune.rds'))

rf_tune <- read_rds(here('code', 'rds', 'rf_tune.rds'))

# extract the best decision model
best_rf <- rf_tune %>%
  select_best("roc_auc")

# print metrics
(rf_metrics <- rf_tune %>%
    collect_metrics() %>%
    semi_join(best_rf) %>%
    select(.metric:.config) %>%
    mutate(model = 'Random forest'))
```

```
## # A tibble: 2 × 7
##   .metric .estimator  mean     n std_err .config              model
##   <chr>   <chr>      <dbl> <int>   <dbl> <chr>                <chr>
## 1 roc_auc hand_till  0.869    10 0.00352 Preprocessor1_Model1 Random forest
## 2 sens    macro      0.643    10 0.00966 Preprocessor1_Model1 Random forest
```

```
# finalize the wf with the best model
rf_workflow <-
  rf_workflow %>%
  finalize_workflow(best_rf)

# generate predictions on the hold-out test data
rf_auc <-
  rf_tune %>%
  collect_predictions(parameters = best_rf) %>%
  roc_curve(.pred_Debris:`.pred_Single worm`, truth = category) %>%
  mutate(model = "Random forest")

rf_auc %>%
  autoplot()
```

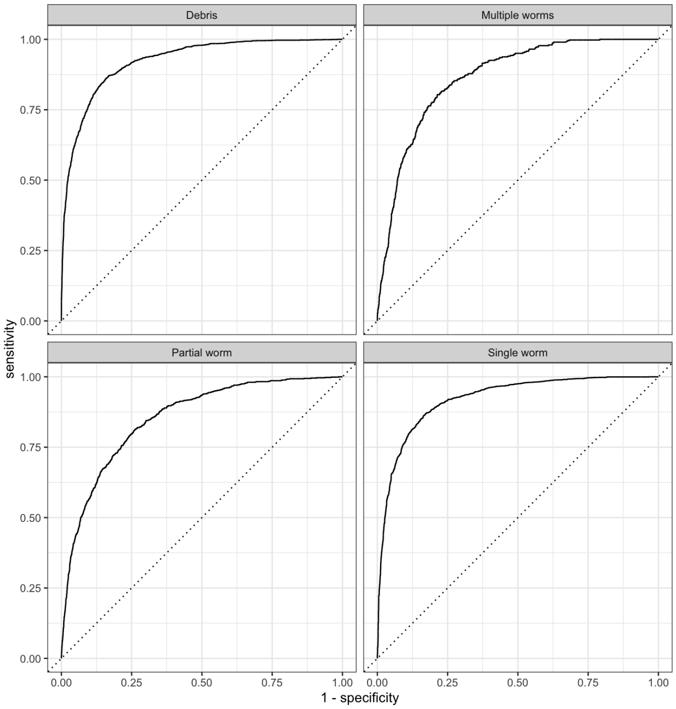


### XGBoost

```r
xg_grid <- grid_latin_hypercube(
  tree_depth(),
  min_n(),
  loss_reduction(),
  sample_size = sample_prop(),
  finalize(mtry(), train_data),
  learn_rate(),
  stop_iter(),
  size = 30
)

# tune on the train data
# xg_tune <-
#   xg_workflow %>%
#   tune_grid(
#     resamples = folds,
#     grid = xg_grid,
#     control = control_grid(save_pred = TRUE,
#                            verbose = TRUE),
#     metrics = metric_set(roc_auc, sens)
#   )
#
# write_rds(xg_tune, here('code', 'rds', 'xg_tune.rds'))

xg_tune <- read_rds(here('code', 'rds', 'xg_tune.rds'))

# extract the best decision tree
best_xg <- xg_tune %>%
  select_best("roc_auc")

# print metrics
(xg_metrics <- xg_tune %>%
    collect_metrics() %>%
    semi_join(best_xg) %>%
    select(.metric:.config) %>%
    mutate(model = 'XGBoost'))
```

```
## # A tibble: 2 × 7
##   .metric .estimator  mean     n std_err .config               model
##   <chr>   <chr>      <dbl> <int>   <dbl> <chr>                 <chr>
## 1 roc_auc hand_till  0.862    10 0.00352 Preprocessor1_Model12 XGBoost
## 2 sens    macro      0.650    10 0.00611 Preprocessor1_Model12 XGBoost
```

```
# finalize the wf with the best tree
xg_workflow <-
  xg_workflow %>%
  finalize_workflow(best_xg)

# generate predictions on the hold-out test data
xg_auc <-
  xg_tune %>%
  collect_predictions(parameters = best_xg) %>%
  roc_curve(.pred_Debris:`.pred_Single worm`, truth = category) %>%
  mutate(model = "XGBoost")

xg_auc %>%
  autoplot()
```

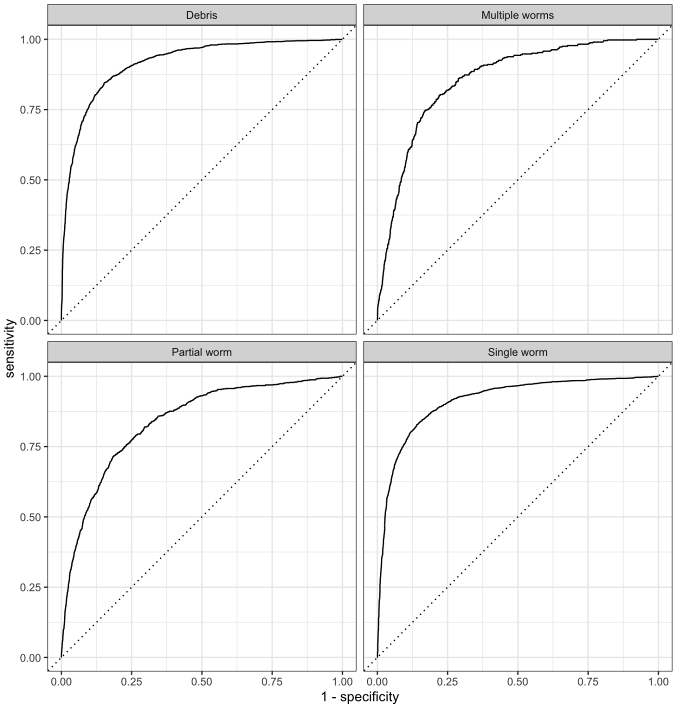


### LightGBM

```r
lgbm_grid <- grid_max_entropy(
  min_n(),
  tree_depth(),
  learn_rate(),
  loss_reduction(),
  size = 30
)

# tune on the train data
# lgbm_tune <-
#   lgbm_workflow %>%
#   tune_grid(
#     resamples = folds,
#     grid = lgbm_grid,
#     control = control_grid(save_pred = TRUE,
#                            verbose = TRUE),
#     metrics = metric_set(roc_auc, sens)
#   )
#
# write_rds(lgbm_tune, here('code', 'rds', 'lgbm_tune.rds'))

lgbm_tune <- read_rds(here('code', 'rds', 'lgbm_tune.rds'))

# extract the best decision tree
best_lgbm <- lgbm_tune %>%
  select_best("roc_auc")

# print metrics
(lgbm_metrics <- lgbm_tune %>%
    collect_metrics() %>%
    semi_join(best_lgbm) %>%
    select(.metric:.config) %>%
    mutate(model = 'LightGBM'))
```

```
## # A tibble: 2 × 7
##   .metric .estimator  mean     n std_err .config              model
##   <chr>   <chr>      <dbl> <int>   <dbl> <chr>                <chr>
## 1 roc_auc hand_till  0.841    10 0.00307 Preprocessor1_Model02 LightGBM
## 2 sens    macro      0.608    10 0.00823 Preprocessor1_Model02 LightGBM
```

```r
# finalize the wf with the best tree
lgbm_workflow <-
  lgbm_workflow %>%
  finalize_workflow(best_lgbm)

# generate predictions on the hold-out test data
lgbm_auc <-
  lgbm_tune %>%
  collect_predictions(parameters = best_lgbm) %>%
  roc_curve(.pred_Debris:`.pred_Single worm`, truth = category) %>%
  mutate(model = "LightGBM")

lgbm_auc %>%
  autoplot()
```

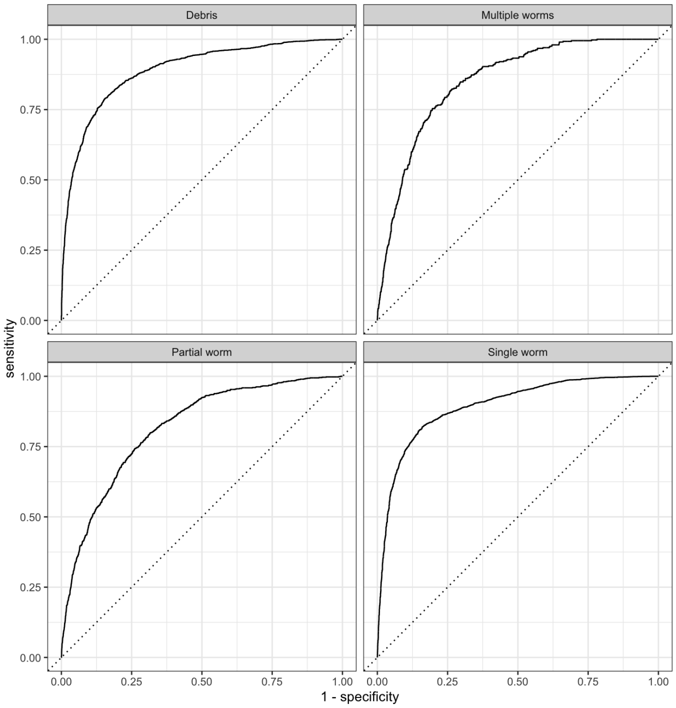


### Evaluate models

```r
(all_metrics <- bind_rows(mn_metrics, rf_metrics, xg_metrics, lgbm_metrics) %>%
   group_by(.metric) %>%
   arrange(-mean))
```

```
## # A tibble: 8 × 7
## # Groups:   .metric [2]
##   .metric .estimator  mean     n std_err .config               model
##   <chr>   <chr>      <dbl> <int>   <dbl> <chr>                 <chr>
## 1 roc_auc hand_till  0.869    10 0.00352 Preprocessor1_Model1  Random forest
## 2 roc_auc hand_till  0.862    10 0.00352 Preprocessor1_Model12 XGBoost
## 3 roc_auc hand_till  0.841    10 0.00307 Preprocessor1_Model02 LightGBM
## 4 roc_auc hand_till  0.802    10 0.00386 Preprocessor1_Model7  Multinomial regr…
## 5 sens    macro      0.650    10 0.00611 Preprocessor1_Model12 XGBoost
## 6 sens    macro      0.643    10 0.00966 Preprocessor1_Model1  Random forest
## 7 sens    macro      0.608    10 0.00823 Preprocessor1_Model02 LightGBM
## 8 sens    macro      0.570    10 0.00746 Preprocessor1_Model7  Multinomial regr…
```

```
(all_models <- bind_rows(mn_auc, rf_auc, xg_auc, lgbm_auc) %>%
  ggplot(aes(x = 1 - specificity, y = sensitivity, col = model)) +
  geom_path(lwd = 1.5, alpha = 0.8) +
  geom_abline(lty = 3) +
  coord_equal() +
  scale_color_viridis_d(option = "plasma") +
  facet_wrap(facets = vars(.level)) +
  theme_minimal() +
  NULL)
```

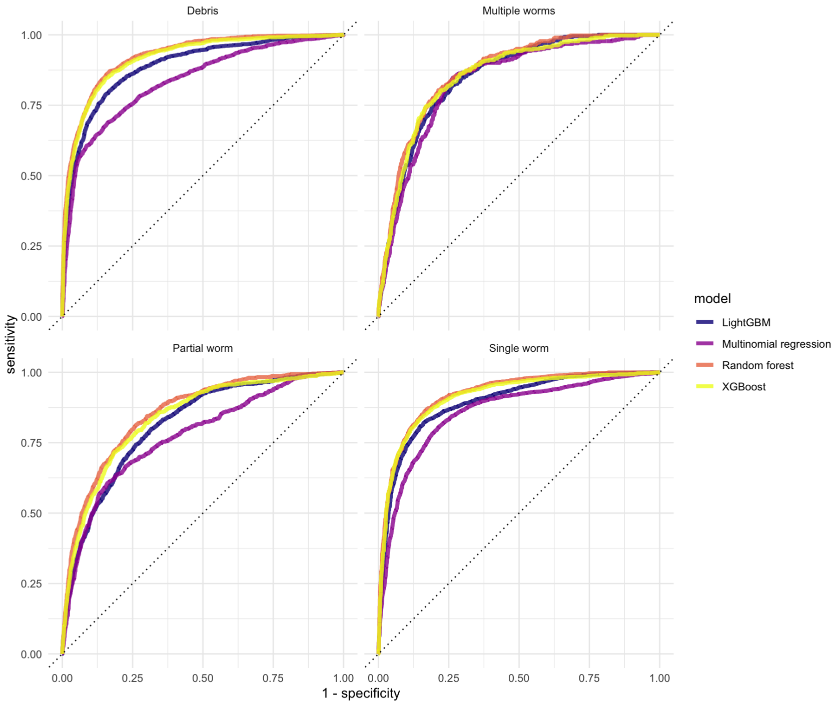


The tree-based methods consistently perform the similarly across all 4 classes. Now fit to the test data using the best parameters and evaluate the model’s performance.

```
mtry <- best_xg$mtry
trees <- 1000
min_n <- best_xg$min_n
tree_depth <- best_xg$tree_depth
learn_rate <- best_xg$learn_rate
loss_reduction <- best_xg$loss_reduction

last_mod <-
  boost_tree(mtry = mtry,
             trees = trees,
             min_n = min_n,
             tree_depth = tree_depth,
             learn_rate = learn_rate,
             loss_reduction = loss_reduction) %>%
  set_engine("xgboost", importance = "impurity") %>%
  set_mode("classification")

last_workflow <-
  xg_workflow %>%
  update_model(last_mod)

set.seed(345)
last_fit <-
  last_workflow %>%
  last_fit(data_split,
           metrics = metric_set(roc_auc, sens))

collect_metrics(last_fit)
```

```
## # A tibble: 2 × 4
##   .metric .estimator .estimate .config
##   <chr>   <chr>          <dbl> <chr>
## 1 sens    macro          0.609 Preprocessor1_Model1
## 2 roc_auc hand_till      0.827 Preprocessor1_Model1
```

```
last_fit %>%
  extract_fit_engine() %>%
  vip::vip() +
  theme_minimal()
```

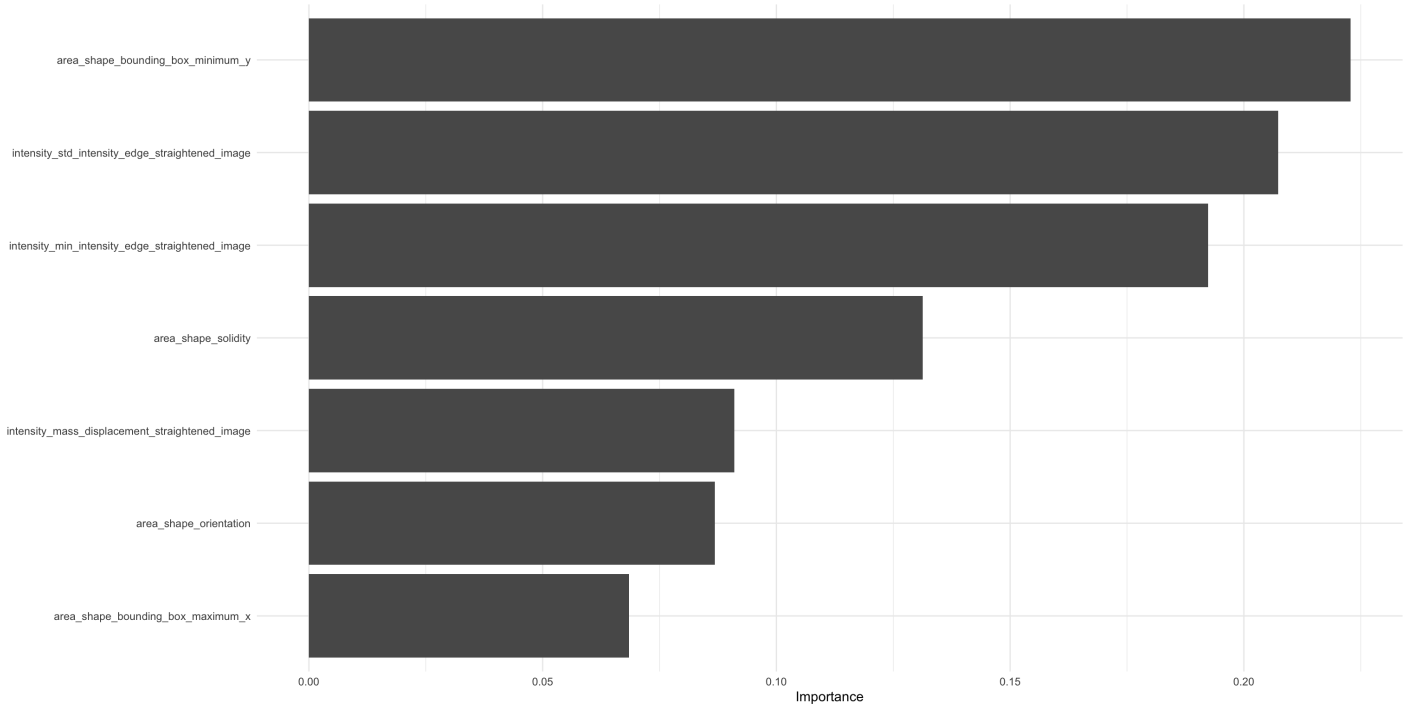


```
(final_auc <-
  last_fit %>%
  collect_predictions() %>%
  roc_curve(.pred_Debris:`.pred_Single worm`, truth = category) %>%
  autoplot())
```

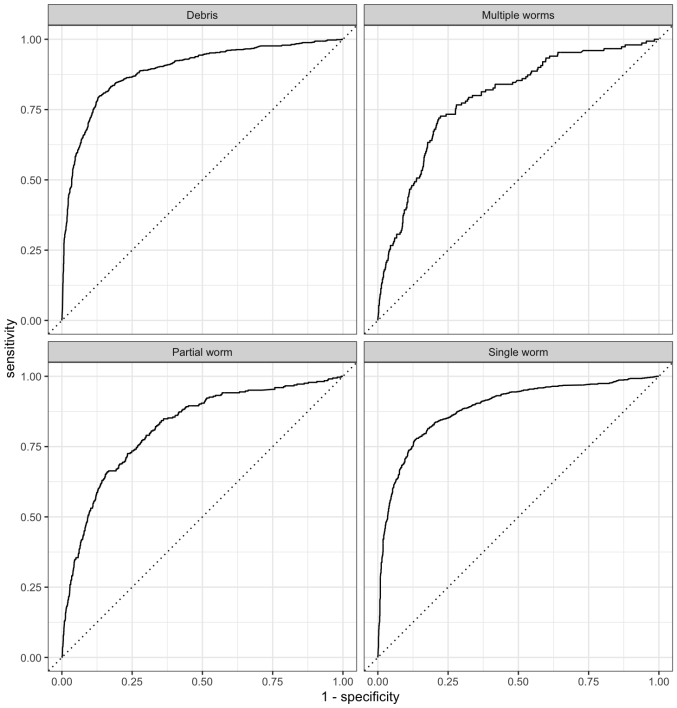


```
last_fit %>%
  collect_predictions() %>%
  conf_mat(truth = category, estimate = .pred_class) %>%
  autoplot()
```

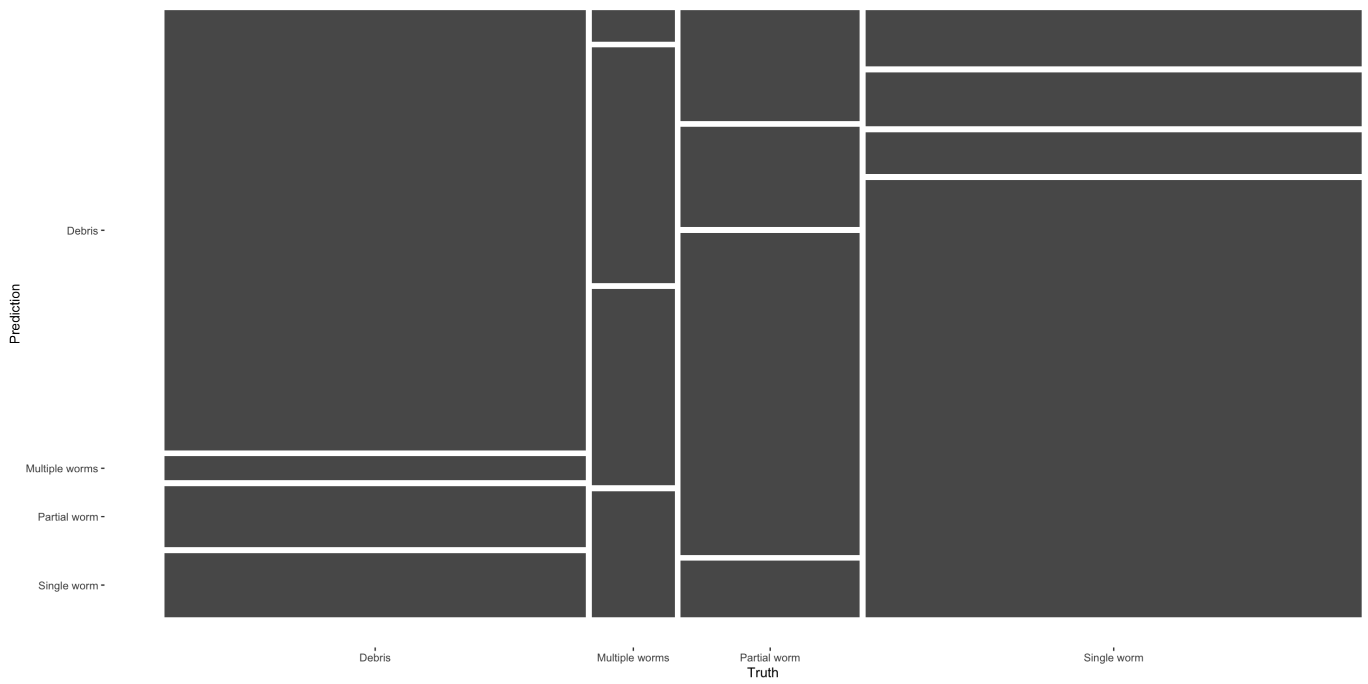


In a situation where we probably have more data points than are truly necessary to be able to draw defensible inferences, we are most concerned with accurate identification of a Single Worm. By that I mean that we are satisfied even if the false negative rate is high (i.e., a Single Worm is identified as either Debris, Partial, or Multiple), as long as those FNs aren't biased towards a certain "type" of worm (i.e., as long as smaller worms aren't more likely to be FNs). Thus, we want a high true positive and low false positive for Single Worms, or high positive predictive value (PPV) and high Sensitivity. Using the selected model and the test set, here's what would happen if we only kept only the StraightenedWorms that were predicted as a Single Worm:

```
last_fit %>%
  collect_predictions() %>%
  filter(.pred_class == 'Single worm') %>%
  conf_mat(truth = category, estimate = .pred_class)
```

```
##                   Truth
## Prediction       Debris Multiple worms Partial worm Single worm
##   Debris              0              0            0           0
##   Multiple worms      0              0            0           0
##   Partial worm        0              0            0           0
##   Single worm        83             32           31         666
```

```
last_fit %>%
  collect_predictions() %>%
  group_by(category) %>%
  summarise(n())
```

```
## # A tibble: 4 × 2
##   category       `n()`
##   <fct>          <int>
## 1 Debris           763
## 2 Multiple worms   150
## 3 Partial worm     324
## 4 Single worm      898
```

```
final_wf <- last_fit %>%
  extract_workflow()

write_rds(final_wf, here('code', 'rds', 'final_workflow.rds'))

pre_filter <- annotations %>%
  select(category, area_shape_major_axis_length) %>%
  ggplot(aes(x = category, y = area_shape_major_axis_length)) +
  geom_quasirandom(aes(color = category)) +
  geom_text(data = . %>% group_by(category) %>% summarise(n = n()),
            aes(label = n), y = 550) +
  theme_minimal() +
  labs(title = 'Pre-filter') +
  lims(y = c(0, 600)) +
  theme(legend.position = 'empty')

post_filter <- augment(final_wf, annotations) %>%
  filter(.pred_class == 'Single worm') %>%
  select(category, area_shape_major_axis_length) %>%
  ggplot(aes(x = category, y = area_shape_major_axis_length)) +
  geom_quasirandom(aes(color = category)) +
  geom_text(data = . %>% group_by(category) %>% summarise(n = n()),
            aes(label = n), y = 550) +
  theme_minimal() +
  labs(title = 'Post-filter') +
  lims(y = c(0, 600)) +
  theme(legend.position = 'empty')

cowplot::plot_grid(pre_filter, post_filter, nrow = 1, align = 'h', axis = 'tb')
```

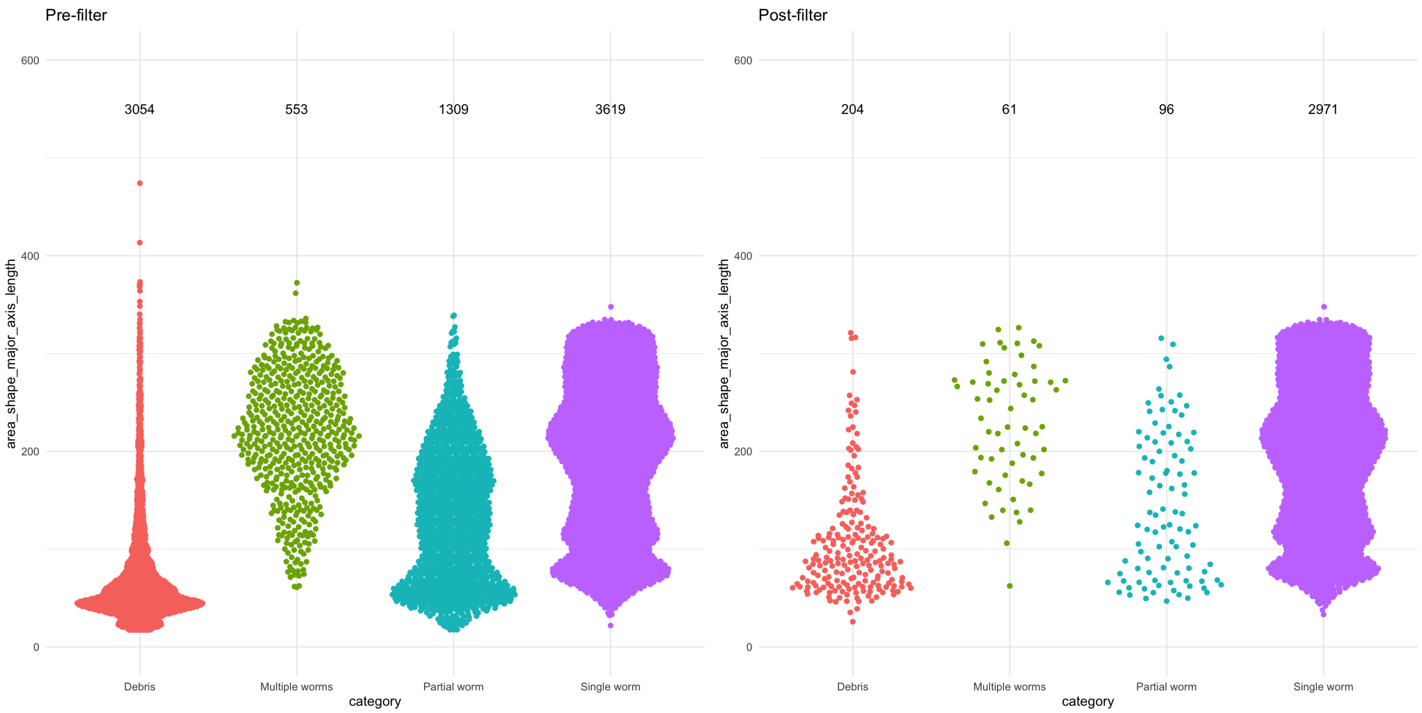


```
(percent_loss <- annotations %>%
  select(category, area_shape_major_axis_length) %>%
  group_by(category) %>%
  summarise(pre_filter = n()) %>%
  left_join(
    augment(final_wf, annotations) %>%
      filter(.pred_class == 'Single worm') %>%
      select(category, area_shape_major_axis_length) %>%
      group_by(category) %>%
      summarise(post_filter = n())
  ) %>%
  mutate(percent_loss = 1 - post_filter / pre_filter))
```

```
## # A tibble: 4 × 4
##   category        pre_filter post_filter percent_loss
##   <fct>                <int>       <int>        <dbl>
## 1 Debris                3054         204        0.933
## 2 Multiple worms         553          61        0.890
## 3 Partial worm          1309          96        0.927
## 4 Single worm           3619        2971        0.179
```
